# Neural resources shift under Methylphenidate: a computational approach to examine anxiety-cognition interplay

**DOI:** 10.1101/2022.04.21.489066

**Authors:** Manish Saggar, Jennifer Bruno, Claudie Gaillard, Leonardo Claudino, Monique Ernst

**Affiliations:** Department of Psychiatry and Behavioral Sciences, Stanford University, Stanford, CA, USA; Section on Neurobiology of Fear and Anxiety, National Institute of Mental Health, Bethesda, MD, USA

## Abstract

The reciprocal interplay between anxiety and cognition is well documented. Anxiety negatively impacts cognition, while cognitive engagement can down-regulate anxiety. The brain mechanisms and dynamics underlying such interplay are not fully understood. To study this question, we experimentally and orthogonally manipulated anxiety (using a threat of shock paradigm) and cognition (using methylphenidate; MPH). The effects of these manipulations on the brain and behavior were evaluated in 50 healthy participants (25 MPH, 25 placebo), using an n-back working memory fMRI task (with low and high load conditions). Behaviorally, improved response accuracy was observed as a main effect of the drug across all conditions. We employed two approaches to understand the neural mechanisms underlying MPH-based cognitive enhancement in safe and threat conditions. First, we performed a hypothesis-driven computational analysis using a mathematical framework to examine how MPH putatively affects cognitive enhancement in the face of induced anxiety across two levels of cognitive load. Second, we performed an exploratory data analysis using Topological Data Analysis (TDA)-based Mapper to examine changes in spatiotemporal brain activity across the entire cortex. Both approaches provided *converging* evidence that MPH facilitated greater differential engagement of neural resources (brain activity) across low and high working memory load conditions. Furthermore, load-based differential management of neural resources reflects enhanced efficiency that is most powerful during higher load and induced anxiety conditions. Overall, our results provide novel insights regarding brain mechanisms that facilitate cognitive enhancement under MPH and, in future research, may be used to help mitigate anxiety-related cognitive underperformance.

## 1. Introduction

Anxiety disorders are highly prevalent in the United States: 19.1% of adults reported having an anxiety disorder in the past year[1], and the lifetime prevalence estimate is 31%[1]. In addition to the severe emotional burden, anxiety interferes with cognition[2] and is associated with cognitive deficits[3–5], further reducing life quality. Paradoxically, performing a task with a high cognitive load can reduce anxiety[6–11]. This latter effect is of particular interest as a potential strategy to optimize anxiety disorders treatment. Studies have sought to understand the impact of pharmacological cognitive enhancement (via Methylphenidate, MPH) on the anxiety-cognition interplay, but the results have been mixed[12, 13]. From a clinical neuroscience perspective, examining anxiety-cognition interactions in the brain is essential to clarify neural mechanisms and lay the groundwork for harnessing this phenomenon for clinical intervention. Here, we use a computational framework that includes both hypothesis-driven and exploratory approaches to advance our understanding of the mechanisms that underlie cognitive enhancement and its impact on anxiety.

Previous theoretical and empirical work has laid a foundation for understanding the interplay between anxiety and cognitive enhancement. First, anxiety (both state and trait) can reduce the i) efficiency of the central executive network; ii) ability to inhibit responses; iii) ability in cognitive switching; and iv) processing efficiency[5, 14–17]. Here, processing efficiency in the context of working memory is defined by the relationship between accuracy and the extent of neural resources mobilized[5]. Thus, high efficiency would indicate higher accuracy while using fewer resources. Second, work in rats has demonstrated that MPH enhances neuronal activity and reduces latency to correct response, showing greater efficiency when performing a visual signal detection task[18]. Finally, neuroimaging during a working memory task has demonstrated that MPH increased activation within the frontoparietal network (FPN) while reducing deactivation within the default mode network (DMN)[19]. This shift in resources was found only during induced anxiety and thus may reflect optimization of the balance between core networks: FPN for regulating cognition[20] and DMN for regulating emotion[21, 22].

While the findings mentioned above present important information regarding how complementary brain networks regulate cognition and anxiety, two critical gaps remain. First, we need to better understand how cognitive load impacts the interplay between induced anxiety and enhanced cognition. Second, there is a lack of understanding about how the whole-brain activity patterns adapt across cognitive load conditions in the presence of induced anxiety or enhanced cognition. The present study gathered data from a randomized trial designed to examine neural mechanisms in response to MPH-related cognitive enhancement and threat-of-shock induced anxiety[19]. Anxiety and cognitive load were experimentally and orthogonally manipulated using threat-of-shock vs. safety and a verbal n-back working memory task at low- and high-load, providing a three-factor design, with the factors being *drug* (MPH vs. placebo), cognitive *load* (1-back vs. 3-back), and *anxiety* (safe vs. induced anxiety) (Fig. 1).

**Fig. 1:**
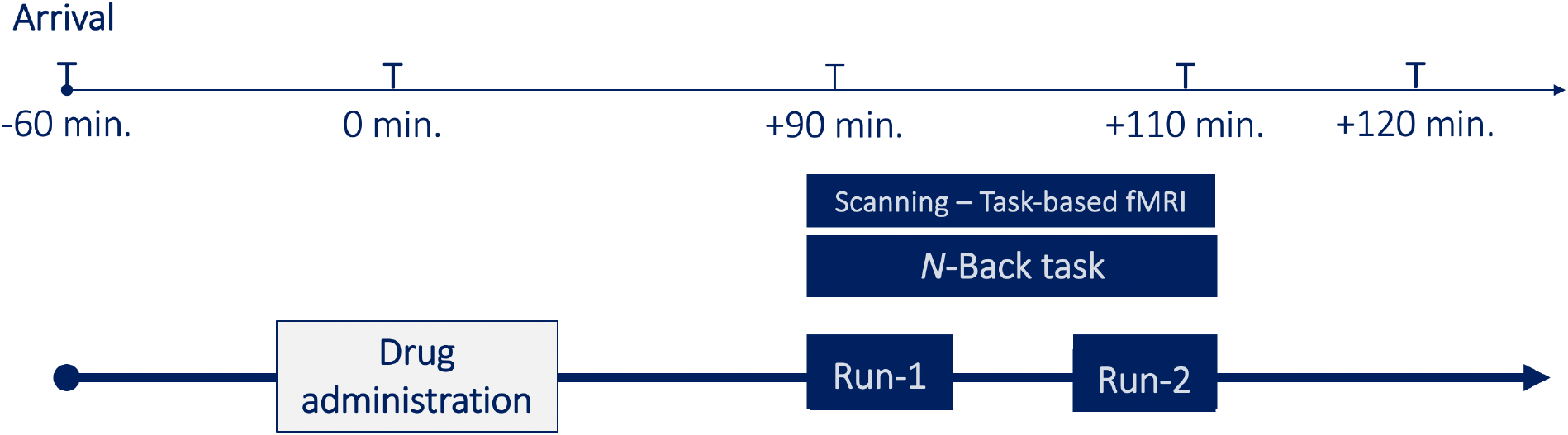
Study timeline for each participant. Drug (Methylphenidate) was administered 90 minutes prior to the beginning of the working memory task in the MR scanner.

We employed complementary hypothesis-driven and exploratory analytical approaches to further understand how boosting cognition via MPH impacts the anxiety/cognitive interplay. These approaches differ fundamentally from a previous study that used a generalized linear model (GLM) to examine changes in brain activity associated with the three-factor design, using the same dataset as here[19]. The hypothesis-driven and exploratory approaches we employed here have unique advantages to the previous GLM-based work. Here, in the first hypothesis-driven computational analysis, we explicitly constructed a mathematical framework to examine how MPH putatively affects cognitive enhancement in the face of induced anxiety across two levels of cognitive load. Hypothesis-driven models typically encapsulate a theoretical (usually mechanistic) understanding of the underlying phenomena and use a comparatively small number of parameters to represent theoretically meaningful constructs[23]. Once fitted to individual data (first-level individual analysis), these parameters can be compared across groups[24]. The hypothesis-driven models are beneficial for measuring hidden variables and their interactions[24], which are otherwise difficult to measure directly. In our hypothesis-driven modeling approach, we defined explicit parameters for cognitive load and anxiety (Fig. 2) to further understand the group differences (MPH vs. placebo) in core network engagement. Based on prior works on the effects of load and anxiety,[19] we limited our examination to the default mode network (DMN) that regulates emotion and the frontoparietal control network (FPN) that regulates cognition.

**Fig. 2:**
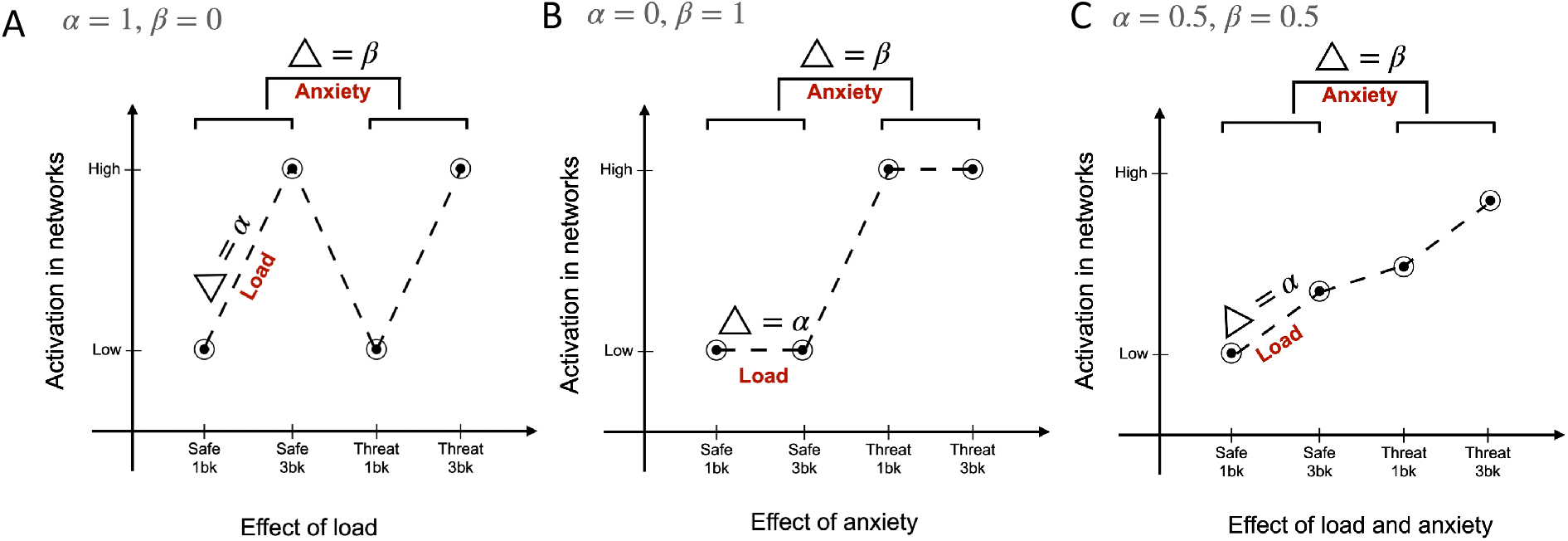
Mathematical formulation and scenarios modeled. We operationalized the framework using two parameters: alpha (*α*) and beta (*β*), where the *α* parameter accounted for the load-related changes in activation (i.e., 3-back > 1-back) and the *β* parameter accounted for the anxiety-related changes (i.e., threat > safe). Using these two parameters, we modeled three different scenarios for each brain network: load-driven, anxiety-driven, and both load and anxiety-driven.

Our second approach used exploratory data analysis to understand how MPH modulates cortical activity patterns at the whole-brain level across varying cognitive load and anxiety. Hypothesis-free (or exploratory) methods designed to detect patterns in whole-brain dynamics without an explicit hypothesis or search restriction have been shown to be essential for understanding the overall dynamic response of the brain to a given task[25]. How the brain dynamically responds to changing cognitive demands is a critical question in clinical neuroscience because aberrant dynamics have been associated with several clinical conditions, including anxiety[26] and ADHD[27]. To examine changes in brain activity patterns without collapsing data in space, time, or across participants at the outset, we used Topological Data Analysis (TDA) based Mapper approach[28]. Mapper attempts to reveal the underlying shape (or manifold) of high-dimensional data by embedding it into a low-dimensional space as a graph. At the same time, the information loss associated with dimensionality reduction is partly salvaged by performing a partial clustering step in the original high-dimensional space[28–30]. We have previously used Mapper to capture task-evoked transitions in whole-brain activity patterns at the acquired spatiotemporal resolution[28, 31], as well as to reveal the rules that govern transitions in spontaneous brain activity at rest[31]. We used whole-brain activity patterns to construct “shape” graphs that portray the manifold governing changes in activity patterns. The nodes in these graphs represent whole-brain configurations, while the edges represent similarity across configurations (Fig. 3). We subsequently annotated (or colored) the graphs based on task conditions and studied differences in topological properties across groups. Together, the hypothesis-driven and exploratory analyses hold the potential for revealing novel, converging evidence to further inform clinical neuroscience.

**Fig. 3:**
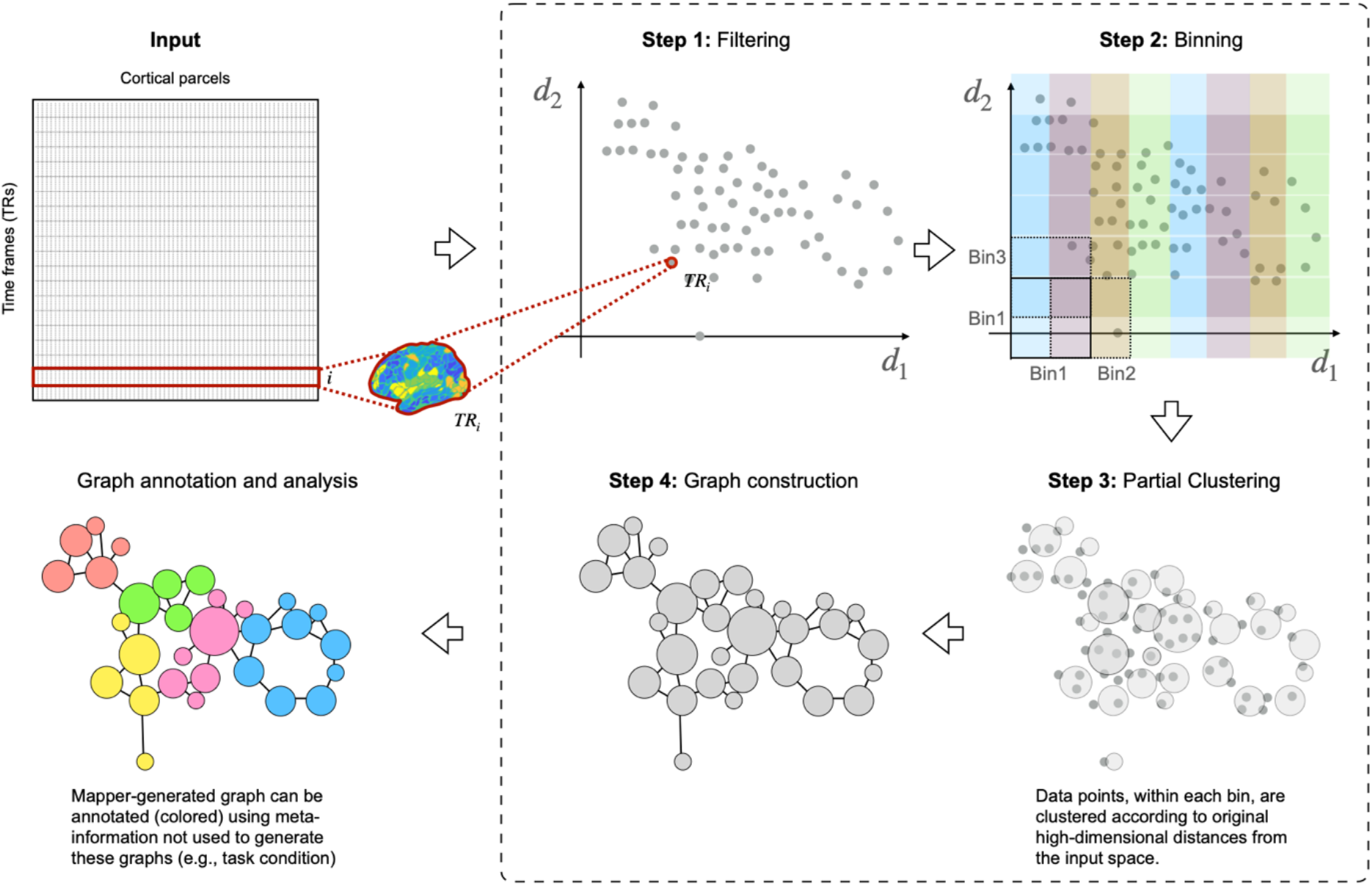
Mapper pipeline is shown pictorially within the dashed border. In the first step, each participant’s high-dimensional neuroimaging data matrix for the entire scan session (time frames x cortical parcels) is embedded into a lower dimension set *d*_*i*_, using a non-linear filter function *f*. In the second step, overlapping *d*-dimensional binning is performed to allow for compression and reduce the effects of noisy data points. In the third step, within each bin, partial clustering is performed such that data points that are closer to each other in the original high-dimensional space coalesce together in the low dimensional space. This partial clustering step allows for recovering information loss incurred due to initial dimensionality reduction. As a final step, to generate a graphical representation of the landscape, nodes from different bins are connected if any data points are shared. Once constructed, Mapper graphs can be annotated (colored) using meta-information, e.g., task condition, and their graph properties can be examined.

## 2. Methods

The present study uses data collected previously as part of the original GLM study[19]. Here we only present brief details about data acquisition.

### 2.1 Participants

Seventy healthy volunteers were recruited from the Washington D.C. metropolitan area in our parallel-group design, a randomized, double-blind placebo-controlled study. Participants were randomly assigned to receive either a single oral dose of 20 mg methylphenidate (MPH) or placebo (PLA), according to a randomization schedule established by the National Institutes of Health (NIH) pharmacy. All subjects provided written informed consent approved by the National Institute of Mental Health (NIMH) Combined Neuroscience Institutional Review Board (CNS IRB). Briefly, participants were aged between 18 and 50 years, with no current psychiatric disorders or past significant psychiatric conditions (assessed by psychiatric interview using the Structured Clinical Interview for DSM-IV (SCID)[32], no medical conditions (assessed by clinical interview and physical exam), and no contraindications to magnetic resonance imaging (MRI). Additionally, participants were excluded if they had prior treatment with stimulants, intelligence quotient (IQ) lower than 80 as assessed via the Wechsler Abbreviated Scale of Intelligence (WASI[33]), pregnancy or a positive pregnancy test, current or past alcohol/drug dependence, alcohol/drug abuse in the past year, or positive toxicology urine screen. Twenty participants were excluded from analyses for the following reasons: missing task-based fMRI sequences (n=4), incomplete behavioral data (n=6), and excessive head motion (n=10). Therefore, the final sample consisted of 50 healthy, right-handed adults (26 F; age mean = 28.2 years, SD = 6.9 years).

### 2.2 Drug administration

Participants received a single oral dose of PLA or immediate-release MPH 20 mg (Ritalin, Novartis, Basel, Switzerland), both presented in identical-appearing capsules. The MPH dose was based on the lowest dose reported to be effective on cognitive function[34–36]. To maximize MPH plasma levels during cognitive testing, the drug was administered approximately 90 minutes prior to the beginning of the experimental working memory task in the scanner (**Fig. 1**)[34, 36, 37]. Potential side effects and adverse reactions were monitored by a clinician using a 34-item inventory to assess physical and mental symptoms (e.g., stomachache, nausea, lightheadedness, rash, drowsiness, headache).

### 2.3 Experimental design

As detailed previously[19], the cognitive task consisted of the widely-used verbal n-back working memory (WM) paradigm[38–40]. Participants viewed letters displayed sequentially on the screen. For each presented letter, participants were instructed to indicate (via button press) if each presented letter was identical to (matched) or different from (did not match); the letter presented *N* letters before. Thirty-three percent of trials were ‘match’ trials in which the presented letter was identical to the letter presented *N* letters before. We included two levels of difficulty: 1-back and 3-back, using a block design. At the beginning of each block, participants viewed instructions (8s) regarding the upcoming difficulty level (1-back or 3-back). The complete task included two runs, each with eight blocks of 18 letters. Each letter was presented for 0.5 s at 2 s intervals. The order of the runs was counterbalanced across participants. Within each run, we also manipulated the level of anxiety by including safe blocks and blocks that included the threat of unpredictable electrical shocks (threat blocks). Each safe and threat block was paired with a level of difficulty (1-back or 3-back). Thus, each run contained two blocks per condition (i.e., 1-back/safe, 3-back/safe, 1-back/threat, 3-back/threat). A color surrounded each letter; either blue to indicate a safe block or orange to indicate a threat block. Participants were explicitly told that they would never receive an electrical shock during the safe blocks (blue) but that they could receive unpredictable electrical shocks at any time during the threat blocks (orange). Three shocks were delivered per run for a total of six shocks throughout the task.

### 2.4 Data acquisition

Two runs of 225 multi-echo EPI images were collected using a 3T Siemens MAGNETOM Skyra (Erlangen, Germany) fMRI system and a 32-channel head coil. Thirty-two interleaved 3mm slices (matrix = 64 mm x 64 mm) were collected parallel to the AC-PC line with an anterior-to-posterior phase encoding direction (TR = 2000 ms; TEs = 12 ms, 24.48 ms, 36.96 ms; flip angle = 70°). Prior to the first functional task-based run, we acquired two additional sets of 10 multi-echo EPI images using the same parameters, with one “forward” series using the same phase-encoding gradient (anterior-to-posterior phase encoding direction) and the second “reverse” series using a reverse phase-encoding gradient with opposite polarity (posterior-to-anterior phase encoding direction). These additional series were used to correct for EPI spatial distortion related to phase-encoding direction. Additionally, a multi-echo T1-weighted Magnetization-Prepared Rapid Gradient-Echo (MPRAGE) image (TR = 2530 ms; TEs = 1.69 ms, 3.55 ms, 5.41 ms, 7.27 ms; flip angle = 7°) was acquired. T1-weighted MPRAGE images consisted of interleaved 1 mm axial slices (matrix = 256 mm x 256 mm), which were later co-registered to the combined EPI images.

### 2.5 Data preprocessing

Results included in this manuscript come from a preprocessing step performed using *fMRIPrep* 1.5.9[41, 42] RRID:SCR_016216), which is based on *Nipype* 1.4.2[43, 44]; RRID:SCR_002502).

#### Anatomical data preprocessing

The T1-weighted (T1w) image was corrected for intensity non-uniformity (INU) with N4BiasFieldCorrection[45], distributed with ANTs 2.2.0RRID:SCR_004757)[46], and used as T1w-reference throughout the workflow. The T1w-reference was then skull-stripped with a *Nipype* implementation of the ants BrainExtraction.sh workflow, using OASIS30ANTs as the target template. Brain tissue segmentation of cerebrospinal fluid (CSF), white matter (WM), and gray matter (GM) were performed on the brain-extracted T1w using fast (FSL 5.0.9, RRID:SCR_002823)[47]. Volume-based spatial normalization to two standard spaces (MNI152NLin6Asym, MNI152NLin2009cAsym) was performed through nonlinear registration with antsRegistration (ANTs 2.2.0), using brain-extracted versions of both T1w reference and the T1w template.

#### Functional data preprocessing

The following preprocessing was performed for each of the 2 BOLD runs per subject. First, a reference volume and its skull-stripped version were generated using a custom methodology of *fMRIPrep*. The BOLD reference was then co-registered to the T1w reference using flirt (FSL 5.0.9)[48] with the boundary-based registration[49] cost function. Co-registration was configured with nine degrees of freedom to account for distortions remaining in the BOLD reference. Head-motion parameters with respect to the BOLD reference (transformation matrices and six corresponding rotation and translation parameters) are estimated before any spatiotemporal filtering using mcflirt (FSL 5.0.9)[50]. The BOLD time series were resampled onto their original native space by applying the transforms to correct for head motion. First, a reference volume and its skull-stripped version were generated using a custom methodology of *fMRIPrep*. Several confounding time series were calculated based on the *preprocessed BOLD*: framewise displacement (FD), DVARS, and three region-wise global signals. FD and DVARS are calculated for each functional run, both using their implementations in *Nipype*[51]. The three global signals are extracted within the CSF, the WM, and the whole-brain masks.

To further reduce the effect of head movement, framewise displacement (FD) was used to create a temporal mask to remove motion-contaminated frames. We used a threshold of FD=0.2mm to flag frames as motion-contaminated. For each such motion-contaminated frame, we also flagged a back and two forward frames as motion contaminated. Following the construction of the temporal mask for censuring, the data were processed with the following steps: (i) demeaning and detrending, (ii) multiple regression using six motion parameters, while temporally masked data were ignored during beta estimation, (iii) interpolation across temporally masked frames using linear estimation of the values at censored frames so that continuous data can be passed through (iv) a band-pass filter (0.009 Hz < f < 0.08 Hz). The temporally masked (or censored) frames were removed for further analysis.

The cortical data was then parcellated into 400 regions using a widely accepted schema that implements a gradient-weighted Markov Random Field (gwMRF) model that integrates both local gradient and global similarity to produce higher functional homogeneity[52].

### 2.5 Hypothesis-based approach: construction of the mathematical framework

We created an explicit mathematical framework to perform a hypothesis-driven examination of the interplay between drug-induced cognitive enhancement and threat-induced anxiety. We operationalized the framework using two parameters: alpha (*α*) and beta (*β*), where the *α* parameter accounted for the load-related changes in activation (i.e., 3-back > 1-back) and the *β* parameter accounted for the anxiety-related changes (i.e., threat > safe). Both parameters were estimated using the BOLD signal strength, 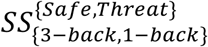, for respective networks *Net*_*<sub>j</sub>*_ and participant *S*_*i*_ as follows,

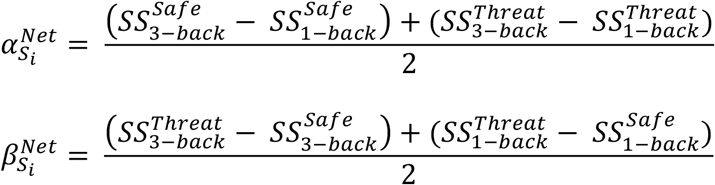

Using these two parameters, for each brain network, we modeled three scenarios: (A) network is only affected by changes in load and not anxiety (i.e., *α* > 0, *β* = 0); (B) network is only affected by changes in anxiety and not load (i.e., *α* = 0, *β* > 0); and (C) network is affected by both load and anxiety (i.e., *α* > 0, *β* > 0). We ignored the null condition, where the network is not affected by either load or anxiety (i.e., *α* = 0, *β* = 0). See **Fig. 2** for a cartoon depicting the parameters and modeled scenarios. Based on previous work, this analysis was limited to default mode network (DMN) and frontoparietal control network (FPN)[19]. The network definitions were based on the Schaefer parcellation[43].

### 2.6 Hypothesis-free approach: TDA-based Mapper pipeline

The TDA[53] based Mapper pipeline was run on each participant. Complete details of the Mapper analysis pipeline are presented elsewhere[28, 29, 31]. The Mapper pipeline consists of four main steps. First, Mapper involves embedding the high-dimensional input data into a lower dimension *d*, using a filter function *f*. For ease of visualization, we chose *d*=2. The choice of filter function dictates which properties of the data are to be preserved in the lower dimensional space. Several studies using animal models and computational research suggest that inter-regional interactions in the brain are multivariate and nonlinear. Thus, we used a nonlinear filter function based on neighborhood embedding[28]. Similar functions have been used previously in manifold learning[54]. Recently, we showed the efficacy of neighborhood embedding (with *k* neighbors) in capturing the landscape of whole-brain configurations extracted from a continuous multitask paradigm and task-evoked data from the Human Connectome Project (HCP)[28].

The second step of Mapper creates overlapping bins in n-dimensional space to allow for compression, thereby reducing the effect of noisy data points. Based on previous work using fMRI data[28], we divided the lower-dimensional space into overlapping bins using a resolution parameter (*r*; #bins) of 20. The percent overlap between bins was kept at 70%. Mapper-generated graphs have been previously shown to be stable for a large variation across parameters for resolution and percent overlap[28, 31]. Here, we varied *k* and *r* across a range of values to examine the robustness of the results (see section 2.8).

The third step of Mapper includes partial clustering within each bin, where the original high-dimensional information is used for coalescing (or separating) data points into nodes in the low-dimensional space. Partial clustering allows recovering the loss of information due to dimensional reduction in step one[29, 55].

Lastly, to generate a graphical representation of the “shape” of input data, nodes from different bins are connected if any data points are shared between them. See **Fig. 3** below for a pictorial representation of the Mapper pipeline.

### 2.7 Network measures

The Mapper-generated graphs can be annotated (or colored) using meta-information not used to construct the graphs. Here, we annotated these graphs using task condition labels to assess whether whole-brain activation patterns are similar or different across task conditions. To quantify the extent of divergence (high condition specificity) or overlap (low condition specificity) across task conditions, we used a graph theoretical measurement of participation coefficient[56]. The participation coefficient of a node is defined as:

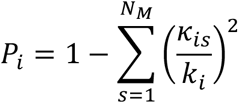

Where *k*_*is*_ is the number of links of node *i* to nodes in community *s, k*_*i*_ is the total degree of node *i* and *N*_*M*_ is the total number of communities. The *P*_*i*_ of a node *i* is close to 1 if its links are uniformly distributed among all communities of the graph (and hence overlapping), and it is close to 0 if its links are primarily within its own community (and therefore diverging).

### 2.8 Effect of Mapper parameters

A parameter perturbation analysis was run to examine whether the results were stable across different parameter choices. Two Mapper parameters, number of neighbors (*k*) and bins (*r*), were varied across a range to show that the observed results using *k*=18 and *r*=20, were robust across other parameter values.

## 3. Results

### 3.1 Behavioral results

Using a linear mixed-effects model, we observed a significant effect of MPH while controlling for age and sex, such that participants in the MPH group outperformed those in the placebo group in terms of accuracy (β = 0.05, t(323) = 1.87, p_(one-tailed)_ = 0.03; we hypothesized that participants in the MPH group would show enhanced performance relative to the PLA thus 1-tailed significance was tested). A more detailed analysis of behavioral results was previously presented[19].

### 3.2 Hypothesis-driven examination using explicit parameters for load- and anxiety-related changes in activation

Based on our previous work[19], we limited the present hypothesis-driven examination of load and anxiety parameters to the default mode (DMN) and frontoparietal (FPN) networks. After estimating load (alpha) and anxiety (beta) parameters for each network, a repeated-measures ANOVA with estimated parameters and networks as within-subject factors and the group as a between-subject factor was run while covarying for age and sex. A significant group x network x parameter interaction was found (F(1,46)=5.335, p = 0.025). Post-hoc pairwise comparisons (adjusted for multiple comparisons using Bonferroni correction) revealed significantly higher values of the load parameter in the MPH group (as compared to PLA) for both networks (p=0.043 for DMN and p=0.009 for FPN). No significant group differences were evident for the anxiety parameter for either network. These results fit with model A (**Fig. 2**). **Fig. 4** shows these results and visualizes each network’s average BOLD signal strength across the two groups and four task blocks.

**Fig. 4:**
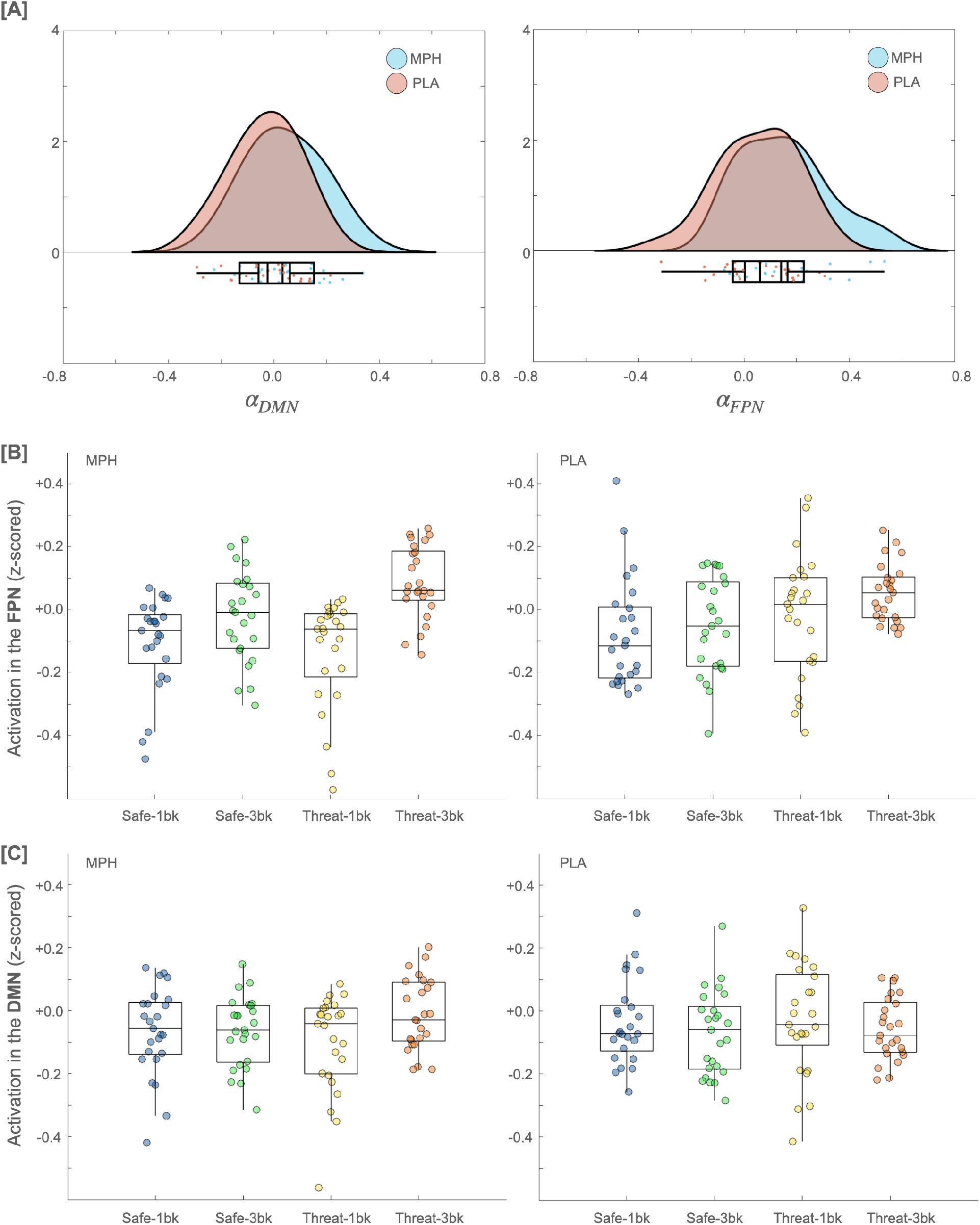
Results of the explicit parameters using default mode network (DMN) and frontoparietal network (FPN). [A] The parameter for the load (alpha) was observed to be significantly **higher** for the MPH group (than PLA group) for both frontoparietal and default mode networks. No significant group difference was observed for the anxiety parameter (beta). [B-C] Shows FPN and DMN networks activation for each of the four conditions across the two groups.

### 3.3 Hypothesis-free examination of induced spatiotemporal changes across the cortex

We used a TDA-based Mapper approach to assess spatiotemporal changes in the entire cortex under different task conditions for the hypothesis-free examination. Mapper graphs were separately generated for each individual using their total task scan (i.e., data combined across both runs). Mapper-generated graphs were later colored (annotated) by load (3-back vs 1-back) and anxiety (safety vs threat) information. See **Fig. 5** for Mapper-generated graphs of representative individuals from both groups. Supplementary Figures **S2-S5** provide Mapper-generated graphs for all individuals, annotated by load and anxiety separately.

**Fig. 5:**
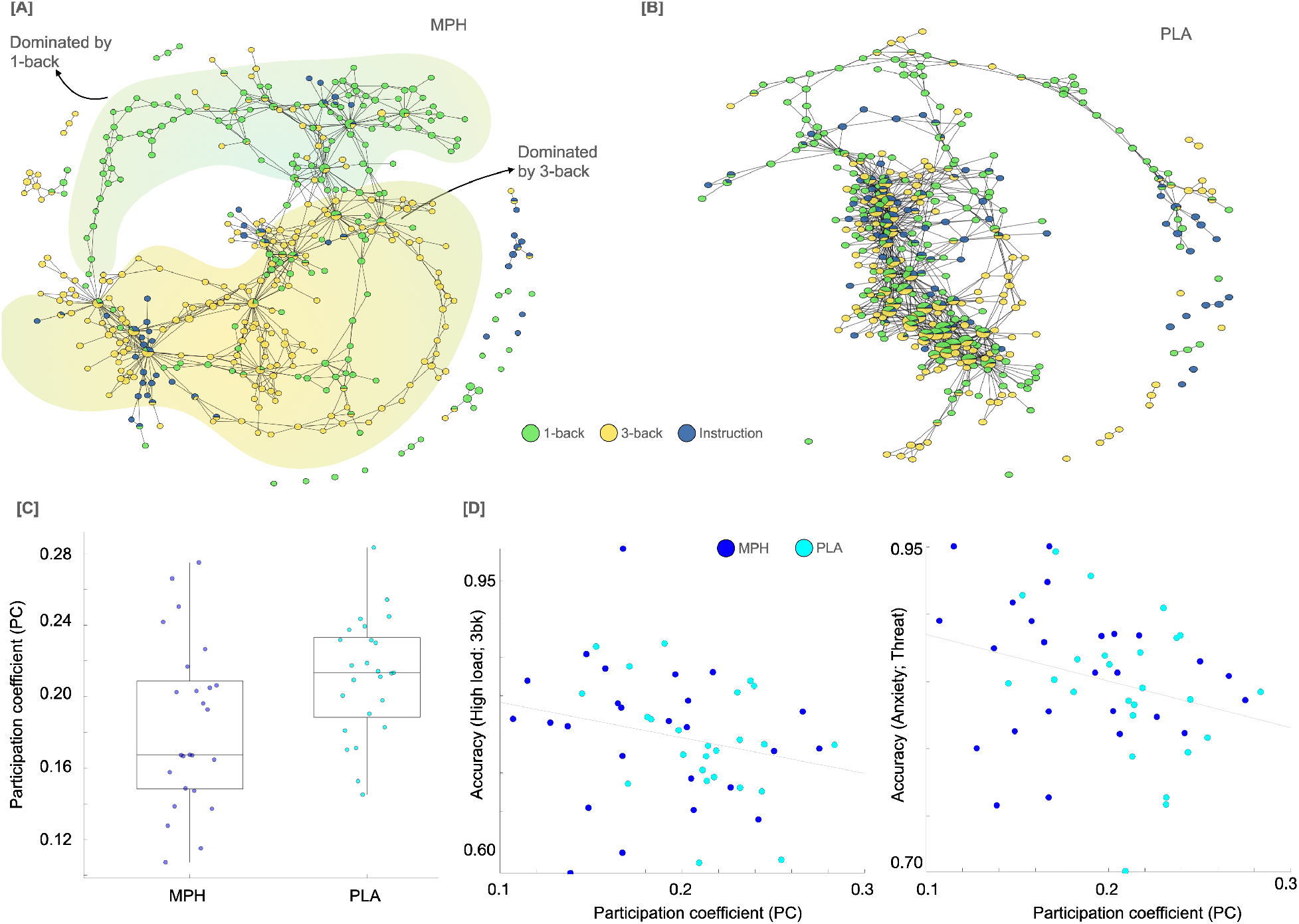
Hypothesis-free examination of induced spatiotemporal changes across the cortex using the topological data analysis (TDA)-based Mapper approach. [A-B] Showing Mapper graphs for two representative participants (methylphenidate [MPH] on the left and placebo [PLA] on the right), annotated by load condition. In the case of the MPH group participant, higher separation between load conditions was observed. To better illustrate the separation between 3-back and 1-back conditions, we added background clusters (yellow and green) to the graph from the MPH participant. [C] The participation coefficient (PC) for load-based annotation was significantly different between the two groups. **[D]** Relating individual differences in participation coefficient of Mapper graphs (extracted from load-based annotation) with behavioral performance on the task, suggesting lower PC values were associated with better performance under both high load (rho (46) = -.31, p=.033) and induced anxiety (rho (46) = -.32, p=.025) conditions.

To estimate the amount of similarity (or divergence) between different degrees of load and anxiety, Mapper-generated graphs colored by load and anxiety were analyzed using the graph theoretical metric of Participation Coefficient (PC)[45]. PC values were then estimated for each graph node. Nodes with higher values of PC indicate higher similarity between different degrees of load (or anxiety), and lower values indicate higher divergence between different degrees of load (or anxiety). A repeated-measures ANOVA with a within-subject factor of annotation (load vs. anxiety) and between-subject factor of the group was run, with age and sex as covariates. A significant group x annotation interaction was found F(1,46)=10.154, p=0.003). Post-hoc pairwise comparisons showed the effect of load to be significantly different across the two groups (p=0.018; Bonferroni corrected), such that significantly lower values of PC were observed for the load-based annotation in the MPH group (as compared with the PLA group). These results suggest higher divergence (or lower similarity) between spatial activity profiles based on the load in the MPH group. No such group differences were found for the anxiety-based annotation of the Mapper graphs. The parameter perturbation analysis showed similar results across different Mapper parameter choices (**Fig. S1**).

Next, we examined whether individual differences in load-related divergence (i.e., lower PC value) were associated with behavioral performance on the working memory task using Spearman’s rank correlation while controlling for age and sex. Data were combined across groups to optimize statistical power. Correlation results indicated that the load-related divergence of Mapper graphs was not only associated with better behavioral performance during higher load (rho (46) = -.31, p=.033) but also during induced anxiety (threat; rho (46) = -.32, p=.025) conditions (**Fig. 5D**).

## 4. Discussion

Using two complementary analytical approaches, we advance our understanding of how cognitive enhancement (under MPH) alters brain activity patterns in the face of induced anxiety and increased cognitive load. In the first approach, we develop an explicit mathematical framework to parametrically investigate the role of two core networks (DMN and FPN) implicated in the anxiety-cognition interplay. In the second approach, we use a hypothesis-free TDA-based analysis to examine the whole-brain dynamical response to our paradigm. Both approaches yield *converging* evidence that cognitive enhancement under MPH facilitates greater differential engagement of neural resources (activation) across conditions of low and high working memory load. This load-based differential management of neural resources facilitated better task performance during both higher load and higher anxiety conditions. Overall, our results provide novel insight into *how* cognitive enhancement under MPH putatively diminishes anxiety. Such information could be critical to informing neurobiologically-targeted treatment approaches employing cognitive enhancement to reduce anxiety and, in turn, reduce anxiety-related cognitive deficits.

Our findings include critical neurobiological and behavioral processes underlying the complex, reciprocal relationship between anxiety and cognition. Previous neuroimaging work has indicated that both DMN, which subserves self-referential and emotional processes[21, 22], and FPN, which subserves executive processes[20], demonstrate aberrant connectivity in association with anxiety disorders[57–60] and with high trait anxiety[60]. Furthermore, emotion regulation is supported by the greater efficiency of both FPN and DMN[61]. Recent neuroimaging work, using the same dataset as here, has shown that cognitive enhancement (via MPH) was associated with increased engagement of the FPN as well as reduced deactivation of the DMN during high anxiety and high WM load condition[19]. This recent finding suggests that expansion of cognitive resources under MPH provides an optimal balance between recruitment of cognitive processing and emotion regulation resources. Using different analytical approaches, we provide complementary evidence suggesting the recruitment of both core networks (DMN and FPN) under MPH.

Additionally, we further specify the nature of the network modulation under MPH according to task difficulty. Specifically, we found that the parameter for the load (model A; Fig 2) was significantly higher in the MPH group relative to the PLA group for both DMN and FPN. Thus, our findings indicate that cognitive enhancement (via MPH) results in a differential engagement in response to a higher working memory load within these networks. Interestingly, no similar group differences were observed for the anxiety parameter, i.e., cognitive enhancement (via MPH) did not result in differential engagement in response to higher anxiety, suggesting a lack of direct interaction between MPH and anxiety processing in our cohort within DMN or FPN.

Our hypothesis-free TDA results provide converging evidence for differential neural engagement in response to higher working memory load under MPH and extend these findings in two important ways. First, the TDA-based Mapper approach is a whole-brain approach and thus provides evidence of whole-brain dynamical response under MPH in addition to the aforementioned network-specific results. Second, our Mapper approach reveals how brain activity patterns differ (i.e., segregate) or collapse (i.e., integrate) at the level of individual timeframes (TRs). We used the participation coefficient[56] (PC), an established graph-theoretical metric, to quantify the degree of segregation (vs. integration) in brain activity patterns across task factors. Using WM load as the task factor revealed lower PC (i.e., higher segregation across WM load) for the MPH group relative to the PLA group. This suggests that cognitive enhancement under MPH was facilitated by differential engagement of neural resources under low and high-load WM conditions. Thus, in line with our network-based approach, the TDA results also suggest load-based differential engagement at the whole-brain level. No such group differences were found for the PC when the Mapper-generated graphs were annotated (colored) by anxiety. Furthermore, WM load-based differential engagement was associated with better behavioral performance during higher load and anxiety conditions. This suggests that by modulating the level of neural engagement, MPH putatively facilitates higher cognitive efficiency during challenging conditions based on required cognitive demand.

Overall, competition for cognitive resources can explain the interactions between anxiety and cognition[62]. When demands for one process increase, the resources available for other processes decrease. The present study was built upon foundational work demonstrating that enhancing cognition with exercise[63] or with MPH[40] results in increased cognitive capacity facilitating enhanced cognitive *and* threat processing. Furthermore, previous work suggested that improving working memory performance through training (practice on high and low load working memory conditions) led to increased anxiety which, as the authors argued, maybe due to the availability of increased resources to process threats[64]. Here, we extend previous work by providing converging evidence that cognitive enhancement associated with MPH is likely facilitated by load-appropriate (and hence efficient) engagement of neural resources. In the future, similar TDA-based analytical approaches can be measured and tracked over time and, as such, may represent a useful metric for informing and tracking response to interventions.

Effective treatment of cognitive interference associated with anxiety disorders is an area of great clinical need. Our study was focused on state anxiety in healthy individuals and thus cannot be generalized to patients with anxiety disorders. The 20mg dose of MPH chosen for this study, while based on previous research[40], was relatively low, yet it resulted in significant neural divergence. This speaks to the sensitivity of our hypothesis-driven and hypothesis-free approaches. However, examining the effects of higher dose MPH will be informative for understanding the whole dynamics of the anxiety/cognition interplay.

In summary, we provide novel mechanistic evidence of load-appropriate engagement of neural resources under MPH. Such efficient load-based engagement was associated with improved behavioral performance in the WM task during high load and anxiety conditions. We hope these results can provide a novel avenue for using computational approaches in improving mechanistic understandings of pharmacological interventions.

## Acknowledgments

M.S. was supported by the NIH Director’s New Innovator Award (DP2; MH119735) and Stanford’s MCHRI Faculty Scholar Award. Financial support for this study was provided by the Intramural Research Program of the National Institutes of Mental Health, project no. ZIAMH002798 to M.E.

## Conflict of Interest

MS: none

JB: none

CG: none

LC: none

ME: none

## Notes

### Competing Interest Statement

The authors have declared no competing interest.

