## Supplementary Information for "Neural resources shift under Methylphenidate: a computational approach to examine anxiety-cognition interplay"

**Running title:** Neural resources shift under Methylphenidate.

*Corresponding authors:
Manish Saggar, Ph.D.,
401 Quarry Rd, St 1356,
Stanford, CA USA 94305
Ph: 650-723-3656


Monique Ernst, M.D., Ph.D.,
15K North Drive
Bethesda MD, 20892
Off: 301-402-9355


[¶] Equal contribution

**Supplementary Information**

### Supplementary Information

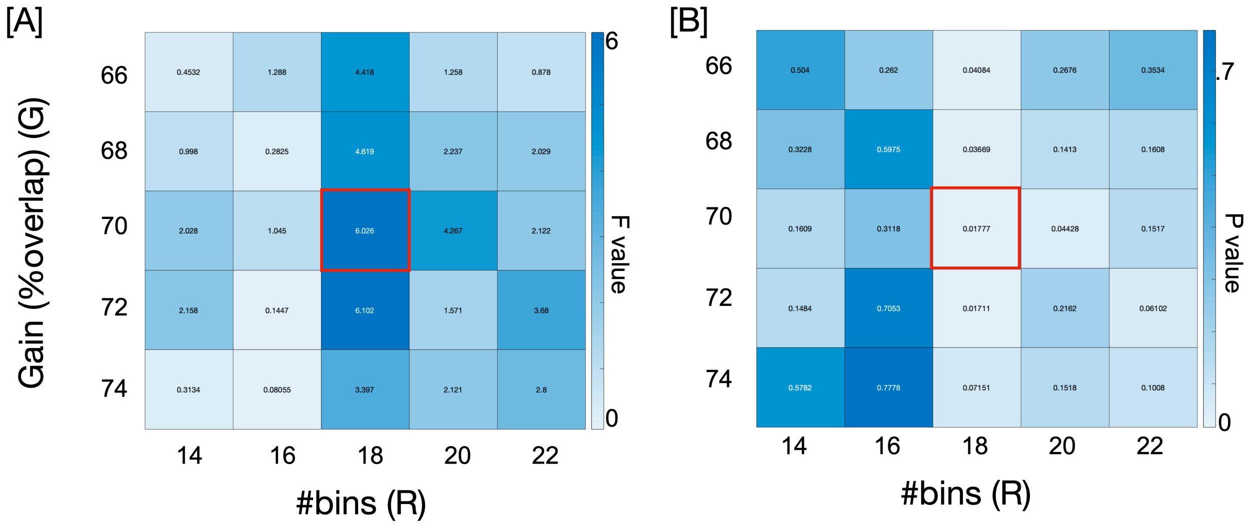


**Fig. S1:** Parameter perturbation for Mapper analysis. We varied two main Mapper parameters Gain (or %overlap between bins) and Resolution (or number of bins) to make sure results were stable across wide parameter choices. The initial values (G=70%, R=18) were chosen based on previous work (ref). As expected, varying both Mapper parameters provide relatively similar results. [A-B] F- and p-values for estimating group differences (from one-way ANOVA) using load-based annotation on the Mapper-generated graphs.

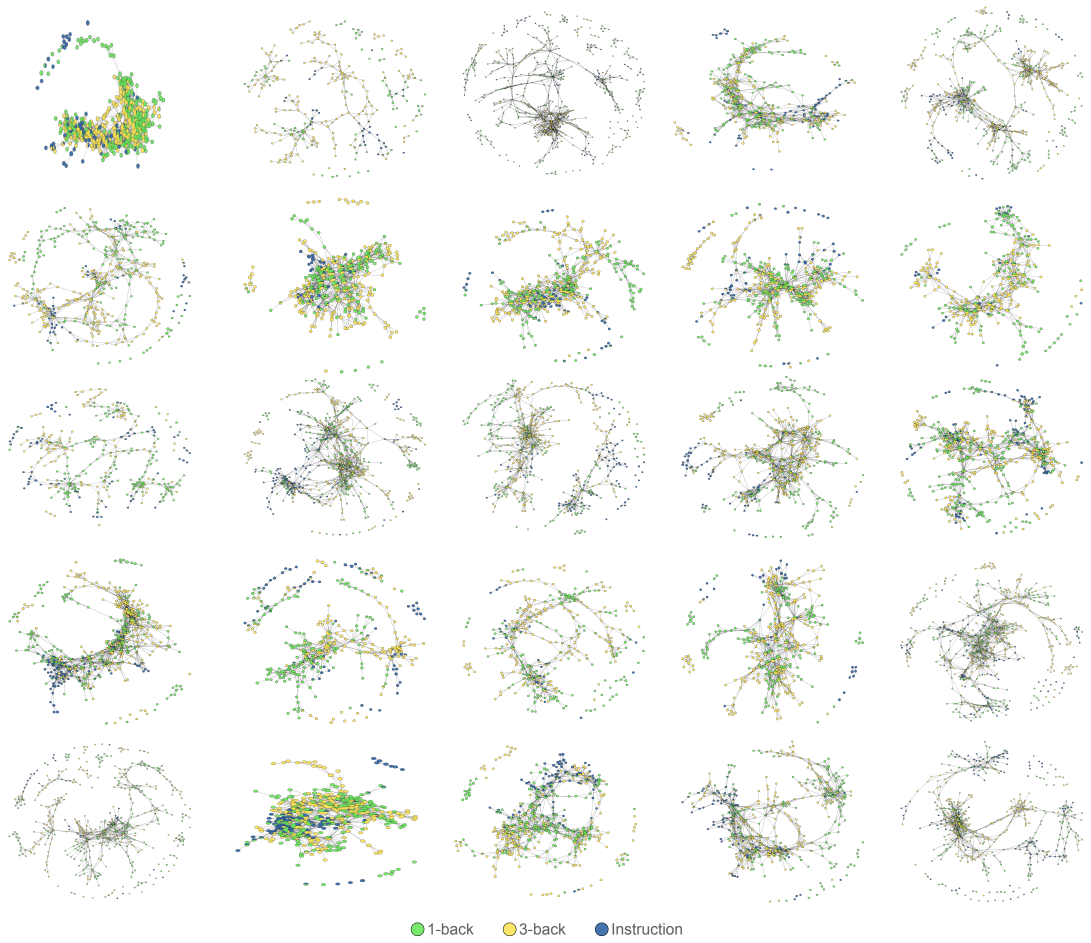


**Fig. S2:** Mapper-generated graphs colored (annotated) by WM load for all participants in the MPH group

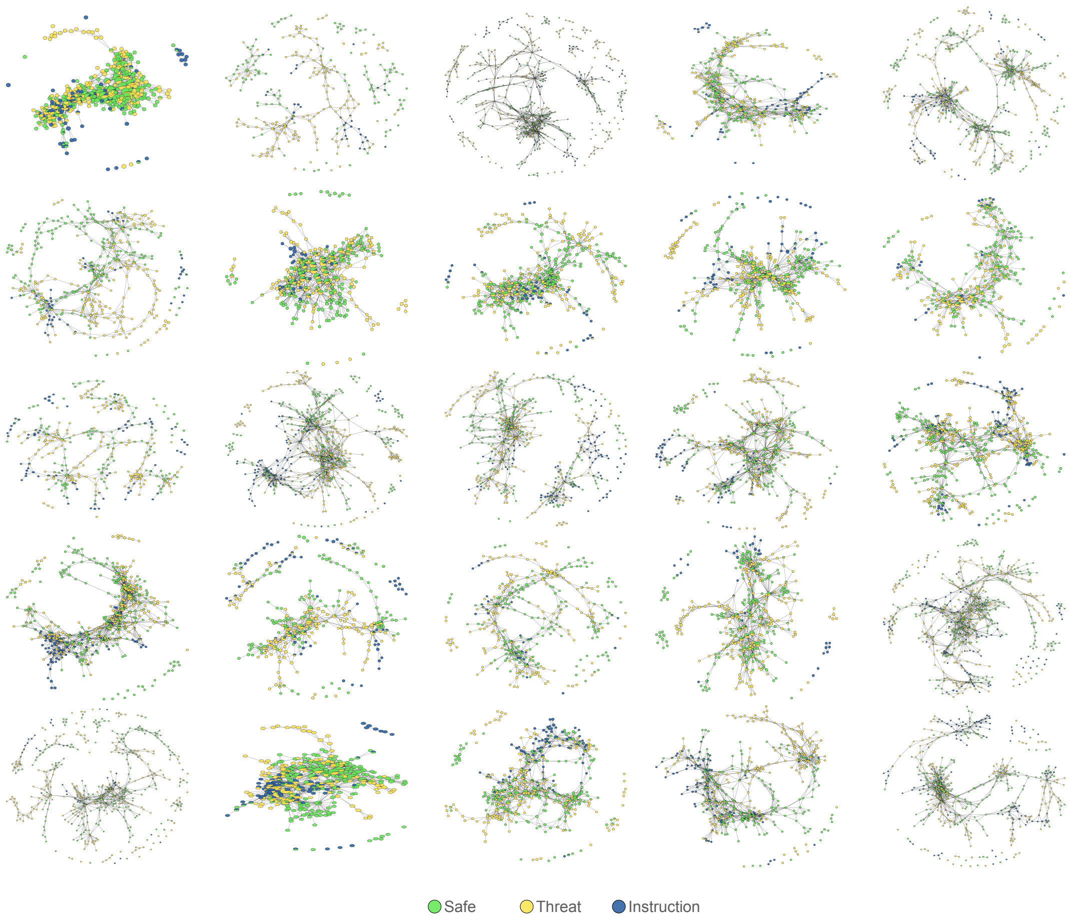


**Fig. S3:** Mapper-generated graphs colored (annotated) by anxiety for all participants in the MPH group

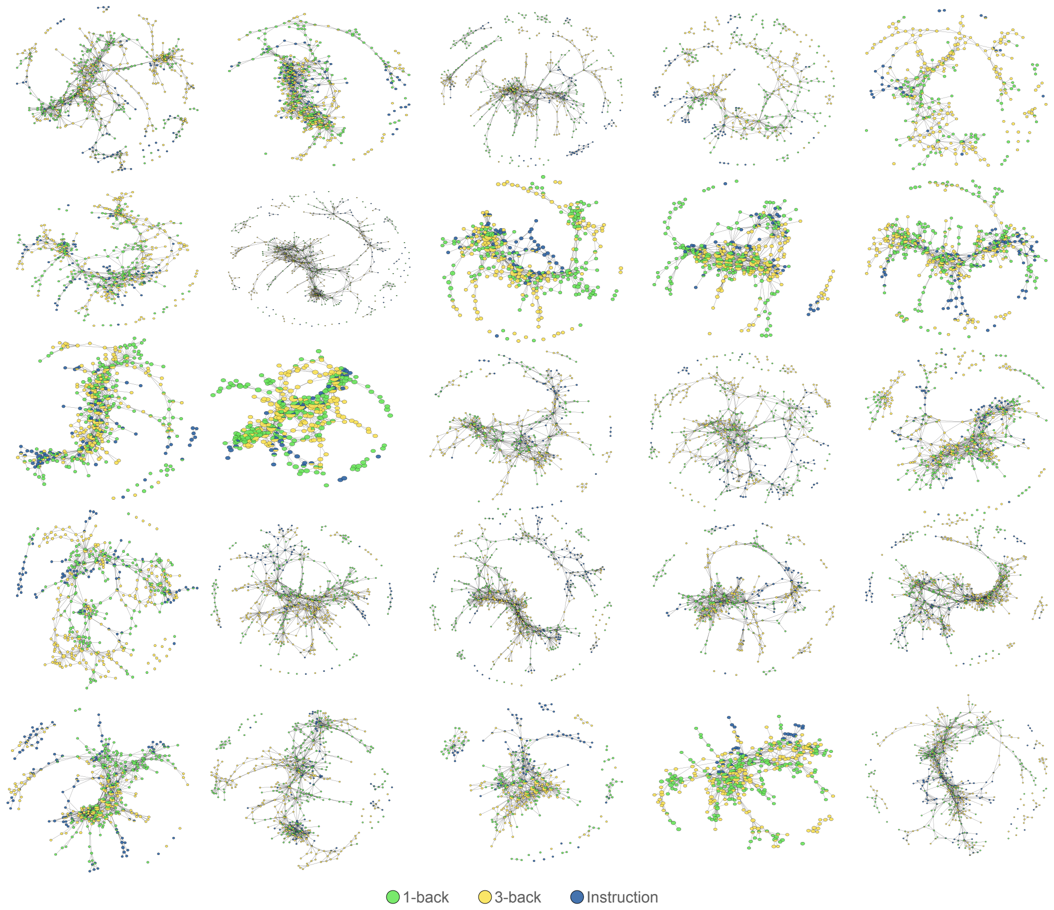


**Fig. S4:** Mapper-generated graphs colored (annotated) by WM load for all participants in the PLA group

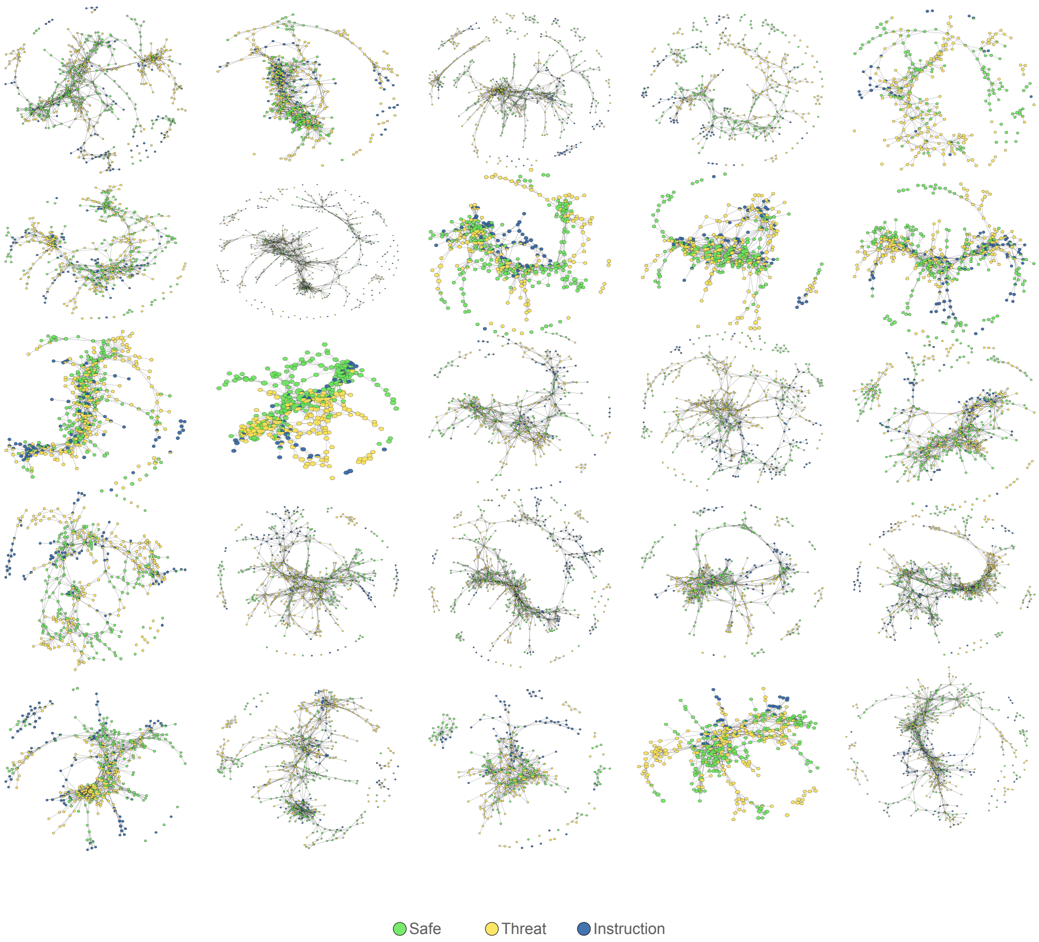


**Fig. S5:** Mapper-generated graphs colored (annotated) by anxiety for all participants in the PLA group
